# Overexploitation counteracts top-down control and the paradox of enrichment in simple food chains

**DOI:** 10.1101/2025.02.26.640097

**Authors:** Josquin Guerber, Nicolas Loeuille, Isabelle Gounand

## Abstract

Overexploitation, the depletion of a resource by its consumer on a short timescale, is widespread in nature but its general implications for biomass control and community stability are not clear. Most approaches investigating the interactions between trophic levels and variations in biomass patterns or in population dynamics generally ignore overexploitation. Here we use a resource-plant-herbivore food chain model allowing for overexploitation (i.e. the plant can overexploit the resource and/or the herbivore can overexploit the plant). We uncover the conditions under which either type of overexploitation occurs and show that they qualitatively change ecological patterns, mainly by suppressing top-down control when interaction strength is high. When plant productivity increases, top-down control patterns are suppressed above the level when the plant starts to overexploit resources. Similarly, when herbivory intensity increases, top-down control patterns disappear when plants become overexploited. Overexploitation also prevents enrichment-driven destabilization by capping the energy fluxes in the community. These findings connect top-down and bottom-up controls in a single framework, and highlight the role overexploitation can play in structuring and stabilizing food chains via the modulation of interaction strengths.

## Introduction

Interactions between primary producers and their consumers are at the base of all natural ecosystems, playing a key role in their maintenance and functioning (Schmitz 2008, Forbes et al. 2019). Since its formulation by Elton in 1927, the concept of food chain has helped to investigate the interactions between trophic levels, highlighting mechanisms that constrain community dynamics and distribution of biomass across trophic levels (Elton 1927). In particular, characterizing the conditions where food chains are under so called “bottom-up” or “top-down” controls, meaning when abundances along the food chain are determined respectively by the abundances of lower or higher trophic levels (Fig. 1), is still an active debate (Barbier and Loreau 2019, Bideault et al. 2021). Classical models highlighting the impact of trophic interactions on population dynamics typically display top-down controls (Oksanen and Oksanen 2000), while models based on the scaling of energy transfer along the food chain imply bottom-up controls (Lindeman 1942). Empirically, top-down and bottom-up controls are typically investigated by exploring the distribution of biomass across trophic levels in natural communities and by manipulating the number of trophic levels in experimental systems. In these settings, for instance, top-down control is associated with inverted biomass pyramids with abundant top consumers and strong trophic cascades, with ongoing debates about their underlying drivers (Chase 2000, McCauley et al. 2018, Su et al. 2021).

**Figure 1:**
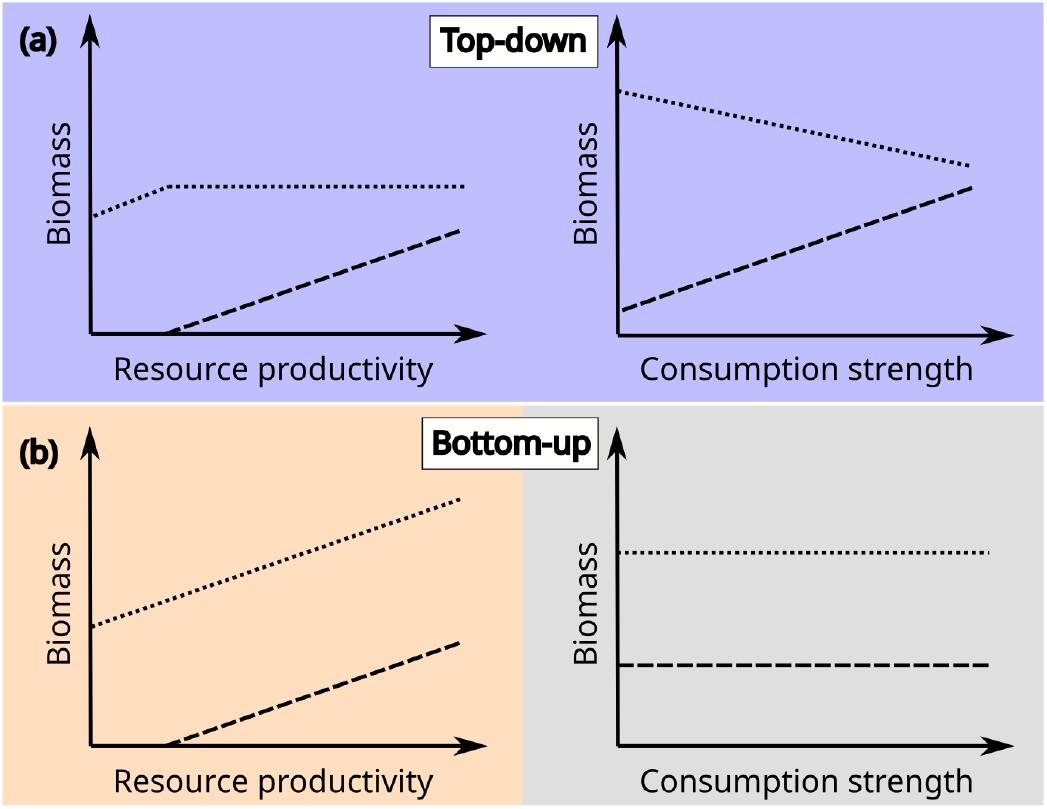
Expected biomass patterns in resource-consumer systems under increase in resource productivity (left) or consumption strength (right). Under top-down control (a), consumer and resource biomass variations (dashed and dotted lines, respectively) are typically anti-correlated. Under bottom-up control (b), consumer and resource biomass share the same variations.

When considering the possibility for control between different trophic levels, it is important to note that a variety of ecological systems exhibit situations where either the nutrient or the primary producer can be almost completely depleted in a very short time (Vuorinen et al. 2021). Because of its high abundance, its high metabolic needs or its behavior, a consumer (plant, herbivore, or higher trophic levels) can completely deplete its resource (nutrient, plant, herbivore or higher) at a shorter time scale than resource regeneration and/or resource population demography. We call this phenomenon “overexploitation”. For example, in a deciduous forest, the outbreak of a folivore insect can completely defoliate trees (Kamata 2002). The mismatch of insect and tree leaves generation times here create an upper limit to plant biomass consumption on a yearly scale. The next year, leaves grow and get consumed again, with both the herbivore and the tree populations being maintained over several years (Kamata 2002). Overexploitation by herbivores is a major issue in both terrestrial (Kallio and Lehtonen 1975, Sharp and Whittaker 2003, Maestre et al. 2022) and aquatic ecosystems (Lampert et al. 1986, Christianen et al. 2014). A trophic level below, primary producers rely on several nutrients and are thus limited by the least available of them (Droop 1968), shaping worldwide patterns of resource limitation (Elser et al. 2007, Browning and Moore 2023). In particular, crops under high agricultural intensity can overexploit soil resources, depleting them more quickly than the resources are able to regenerate (Tan et al. 2005).

The possibility for overexploitation is often overlooked when investigating top-down or bottom-up regulations. Classical consumer-resource dynamical models based on Lotka-Volterra (Lotka 1925, Volterra 1931) or Rosenzweig-MacArthur approaches (Rosenzweig and MacArthur 1963) emphasize the impact of consumers on resources rather than on resource availability. In these models, a higher consumer abundance therefore always increases consumption and the limitation by the resource is either implicit (through the use of a carrying capacity) or an emergent property (through changes in resource abundance). This may lead to overshooting effects due to delays between the growths of consumers and resources, where the resource can be completely depleted, leading to the crash of the system (Rosenzweig 1971). Conversely, models based on the scaling of energy transfer between trophic levels imply bottom-up control, as the energy available to a trophic level cannot exceed the energy represented by the biomass of the lower trophic level, which therefore conditions its existence (Lindeman 1942). However, these energy-based models mainly describe steady states and do not investigate explicit population dynamics. Others have introduced dynamical models that allow to focus on the lower trophic levels’ availability. In Droop’s model, the growth rate of an algae population is determined by the intracellular concentration of its limiting nutrient (Droop 1968). With ratio-dependent functional responses (Arditi and Ginzburg 1989), resource limitation depends on both species net and relative abundances, which prevents overshooting effects and exhibits bottom-up control patterns. Similarly, functional responses that saturate at high resource abundance (e.g., Holling type II or type III) limit overshooting effects by capping interaction rates. However, these models do not take into account a possibility of persistent resource depletion that would emerge from a difference in time scales between consumption and resource regeneration.

Dynamical consumer-resource models on one side and energy-based, resource-focused models on the other side have provided different insights on producer-consumer interactions, that may or may not hold when accounting for overexploitation (Fig. 1). Systems with a heavy top-down control imply that populations may only be capped by the presence of their consumers (L. Oksanen et al. 1981). In such models, increasing plant productivity in a plant-herbivore system (without upper levels) increases herbivore biomass while the plant stays at the same biomass (Fig. 1a), given the top-down control exerted by herbivores (Oksanen and Oksanen 2000). When ecosystem productivity allows the persistence of a carnivore which exerts a top-down control on the herbivore, herbivore biomass stops increasing or even decreases, thereby enabling plant abundance to grow (i.e., a trophic cascade, Fig. 1a). In contrast, in systems with strong bottom-up control, higher basal productivity increases the biomass of all trophic levels (Frederiksen et al. 2006), while productivity increases in higher trophic levels do not affect biomass stocks at lower levels (Fig. 1b). Variations in biomass with productivity of adjacent trophic levels are thus positively correlated under bottom-up control, and anti-correlated under top-down control (Fig. 1). Reconciliation of the two paradigms is emerging in ecological theory (Barbier and Loreau 2019, Galiana et al. 2021), showing the strength of trophic interactions as a key parameter for determining bottom-up and top-down control. When interaction strength scales with consumer metabolism, the control is more top-down (Barbier and Loreau 2019). Conversely, when interaction strength scales with resource metabolism, bottom-up control dominates (Barbier and Loreau 2019). We argue that while these models nicely represent exploitation dynamics in a number of situations, they do not capture over-exploitation effects when resource regeneration and consumer demography act on decoupled time scales. Implementing explicit overexploitation in consumer-resource models could thus question the conditions under which bottom-up or top-down controls occur and how they shape biomass patterns along productivity gradients.

Overexploitation also likely constrains the stability of ecological dynamics. As suggested by Rosenzweig in 1971, the paradox of enrichment states that energy or resource input in simple food chains can destabilize consumer-resource interactions, inducing oscillations or even the crash of the system. Later works showed that strong interaction strength destabilizes communities (McCann et al. 1998), helping to build a generic theory that states that strong energy fluxes destabilize food chains (Rip and McCann 2011). Indeed, enriched systems exhibit higher energy fluxes and are therefore less stable. However, strong trophic interactions could also favor resource overexploitation, which could potentially affect stability, although this aspect is not explicitly considered in these models.

The role of possible overexploitation for shaping the direction of ecological control as well as biomass patterns and dynamical regimes still needs to be investigated. We here build a simple food chain model with recycling, integrating overexploitation of resources by plants (thereafter “resource overexploitation”), of plants by herbivores (thereafter “plant overexploitation”), or both at the same time. With this model, we question how overexploitation alters plant-herbivore interactions by answering three main points: (i) The conditions under which coexistence and overexploitation of either type occur. We expect coexistence when both plant and herbivore productivities are sufficient relative to their mortality, with overexploitation occuring when consumption becomes too high. (ii) The modifications of classical patterns linking productivity and standing biomass given overexploitation. We expect that plant overex-ploitation can cap herbivore biomass, even in the absence of carnivores and contrary to classical dynamical consumer-resource models. However, we also expect that, without overexploitation, increasing herbivore consumption should decrease or cap plant biomass, due to top-down control. (iii) The effects of overexploitation on stability. We expect that while stronger interactions and enrichment decrease the stability of the system, this effect can be limited by overexploitation. Our results show that overexploitation, favored by stronger interactions, introduces bottom-up control in an otherwise top-down controlled system, suppressing classically expected biomass and stability patterns.

## Methods

We built a discrete-time plant-herbivore food chain model with nutrient recycling and explicit overexploitation possibility for both nutrient and plant by their respective consumer. We analytically calculated steady states and analyzed how their biomass composition and stability varied with the consumption parameters: plant productivity and herbivory intensity. Varying these parameters allows us to highlight conditions leading to overexploitation and the associated dynamical regimes.

### Model description

We express the dynamics of the mass of a single nutrient as it is consumed and recycled through three compartments: inorganic resources *R*, plants *P*, and herbivores *H*. We use a discrete time scale *t* where a time step represents the time scale on which overexploitation may occur: when the mass of resource required by a consumer for its growth between *t* and *t* + 1 exceeds the available mass of its resource at time *t*, the resource is overexploited. We assume that the total (bio-)mass of a compartment is proportional to the mass of nutrients it contains. The dynamics within the food chain are determined by the following system:

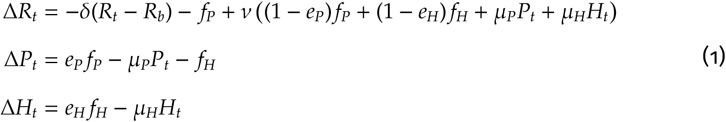

These equations model four phenomena:

- Basal resource dynamics as a chemostat tending towards resource mass *R*_*b*_ given a diffusion rate δ.
- Feeding intensity of plants on resources *f*_*P*_ and of herbivores on plants *f*_*H*_ (see below for the detail of their expressions), with conversion efficiencies *e*_*P*_ and *e*_*H*_.
- Metabolic losses for the plant and herbivore compartments, µ_*P*_ and µ_*H*_.
- Recycling proportion ν of the loss terms from the biotic compartments to the resource compartment. Nutrient mass that is not recycled is lost by the system (e.g. by leaking to surrounding ecosystems).

Feeding terms for the plant and the herbivore compartments are defined by the following functions :

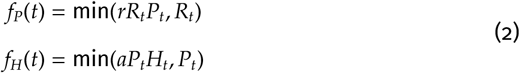

These minimum functions allow us to distinguish two regimes: (i) available resource is sufficient and feeding is a Lotka-Volterra linear term, where the consumption of the resource depends both on resource and consumer masses and (ii) available resource would not be sufficient to cover the Lotka-Volterra interaction term, so that all the resource is consumed (overexploitation). Plant and herbivore feeding are scaled using the parameters *r* (hereafter “plant productivity”) and *a* (hereafter “herbivory intensity”), respectively. Note that, while system (1) could theoretically lead to negative biomass stocks, we restricted our analyses to parameter values that never yielded negative biomasses during simulations.

All symbols, their definitions, and their default values are summarized in Tab. 1.

**Table 1:**
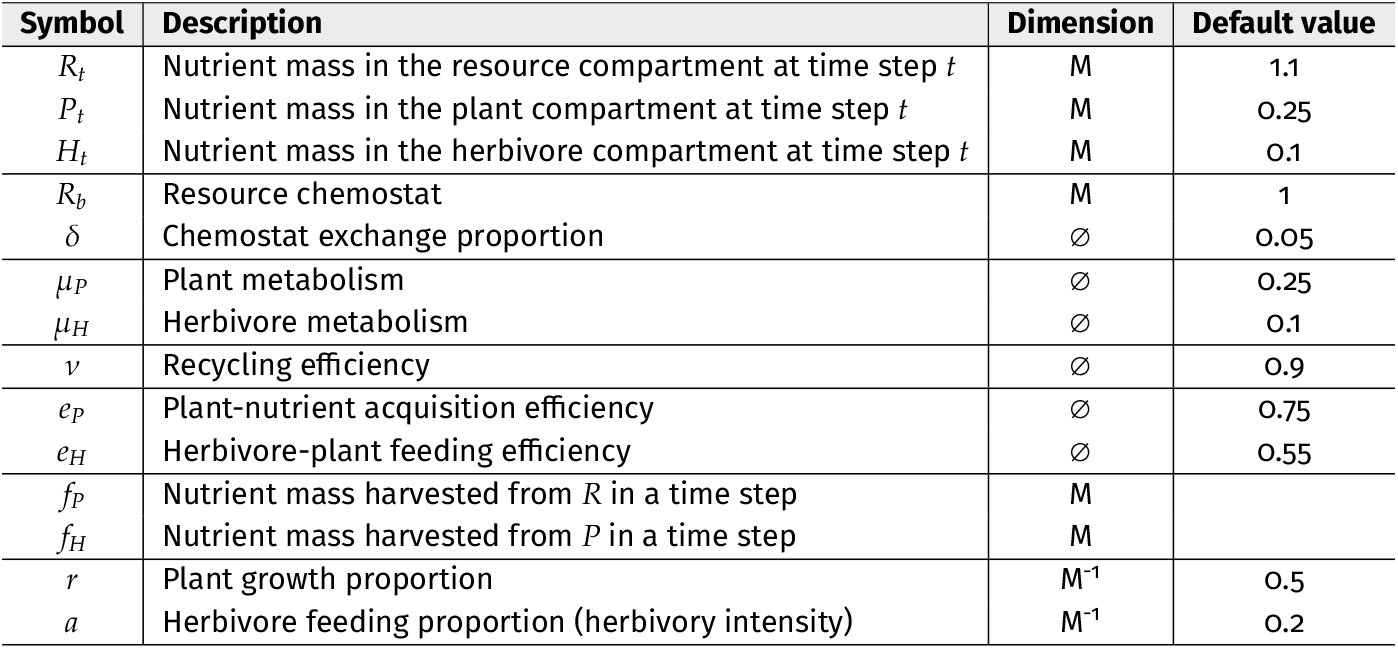
Symbols used in equations and figures. For each symbol, we provide a short description, its dimension and the default value used in most simulations. Values for *R*_*t*_, *P*_*t*_ and *H*_*t*_ are used as initial conditions in numerical simulations.

### Analytical and numerical methods for the analysis of the model

Depending on functions *f*_*P*_ and *f*_*H*_, the complete system can yield 4 different outcomes. Eq. 2 implies that resource and plant overexploitations occur when *P*_*t*_ > 1/*r* and *H*_*t*_ > 1/*a*, respectively. Based on these inequalities, 4 situations may occur: no overexploitation, overexploitation of resources, of plants, and overexploitation of both resources and plants. For each situation, we analytically obtained explicit expressions for each compartment’s mass at steady state (fixed points). The expressions found for the fixed points of a given situation are restrained by the overexploitation conditions of the concerned situation, yielding *validity* boundaries between situations. We estimated the stability of fixed points by numerically computing the dominant eigenvalue of the Jacobian of the system around each fixed point. This allowed us to distinguish whether each fixed point corresponded to a stable equilibrium, eventually reached by damped oscillations (stable node or stable spiral), or if it was unstable (Murray 2002). We characterized the dynamics around unstable states (e.g. presence of cycles) by inspecting simulations. For each parameter combination, we thus either could identify a single valid and stable fixed point or noticed that the system produced cycles around the values in Equation S7.

We investigated the influence of plant productivity *r* and herbivory intensity *a* on the system biomass patterns and stability by combining analytical results with simulations of system dynamics. When deriving the analytical expression for each fixed point with respect to *r* (respectively to *a*), the sign of the derivative directly gives the sign of the variation in compartment mass under an increase of *r* (respectively *a*). We further illustrate the results from the derivatives by plotting compartment mass patterns under variations of *r* or *a*. To do so, we examined the values of *R, P* and *H* over the last third of simulations to characterize long-term dynamics. We (i) distinguished which overexploitation situation each parameter combination corresponded to, by averaging the biomass values and comparing them to the analytical expressions for the fixed points and (ii) checked the presence of oscillations (even damped), by detecting the local extrema for each compartment. Simulations were run over 500 time steps, which was sufficient to reach steady states, using the Julia programming language (Bezanson et al. 2012), with the help of the DynamicalSystems.jl package (Datseris 2018). See (Fig. S1) for examples of simulation outputs.

To test our prediction that overexploitation should introduce an upper cap in the increase of herbivore biomass with plant productivity *r*, we explored the effects of varying *r* from close to 0 to 10 for two different values of herbivory intensity. To test that increasing herbivory intensity *a* should decrease the stability of the system (Rip and McCann 2011) and favor plant overexploitation, we explored *a* from close to 0 to 10 for three different values of plant productivity (see Fig. S2 for a representation of explored parameter ranges as transects across the parameter space). Larger ranges for *r* and *a* did not lead to regimes that qualitatively differed from those presented here. Values for the other parameters were set according to Tab. 1: δ reflected a slow chemostat and ν an efficient recycling. µ_*P*_ and *r* were set higher than µ_*H*_ and *a*, respectively, to ensure that plant population dynamics could be maintained both with and without herbivores. Herbivore trophic efficiency was set as *e*_*H*_ = 0.55, following Yodzis and Innes 1992. We checked that our results did not substantially change for lower recycling efficiencies (Fig. S3).

## Results

Community composition and overexploitation patterns interact in complex ways. Sufficient plant productivity, *r*, and herbivory intensity, *a*, allow coexistence of resources, plants and herbivores (Fig. 2a, light gray area). Further increasing *r* or *a* leads to over-exploitation, with overexploitation of both resources and plants occurring at high plant productivity and high herbivory intensity (Fig. 2b, blue area). The intersection between coexistence and overexploitation conditions yields seven different situations (Fig. 2c). When plants are alone, they can overexploit resources only when their productivity is sufficiently high (regime B1 in Fig. 2c). Also, note that the top-down control exerted by herbivores is not always sufficient to prevent overexploitation of resources by plants: resource, plant, both or no overexploitation are all possible when the three compartments co-exist. Analysis of stability shows that most equilibria are stable. Limit cycles however occur at low productivity and high herbivory in the regime without overexploitation (D3 in Fig. 2c). Including this unstable outcome therefore yields an eighth possible situation, highlighted in brown on Fig. 2c.

**Figure 2:**
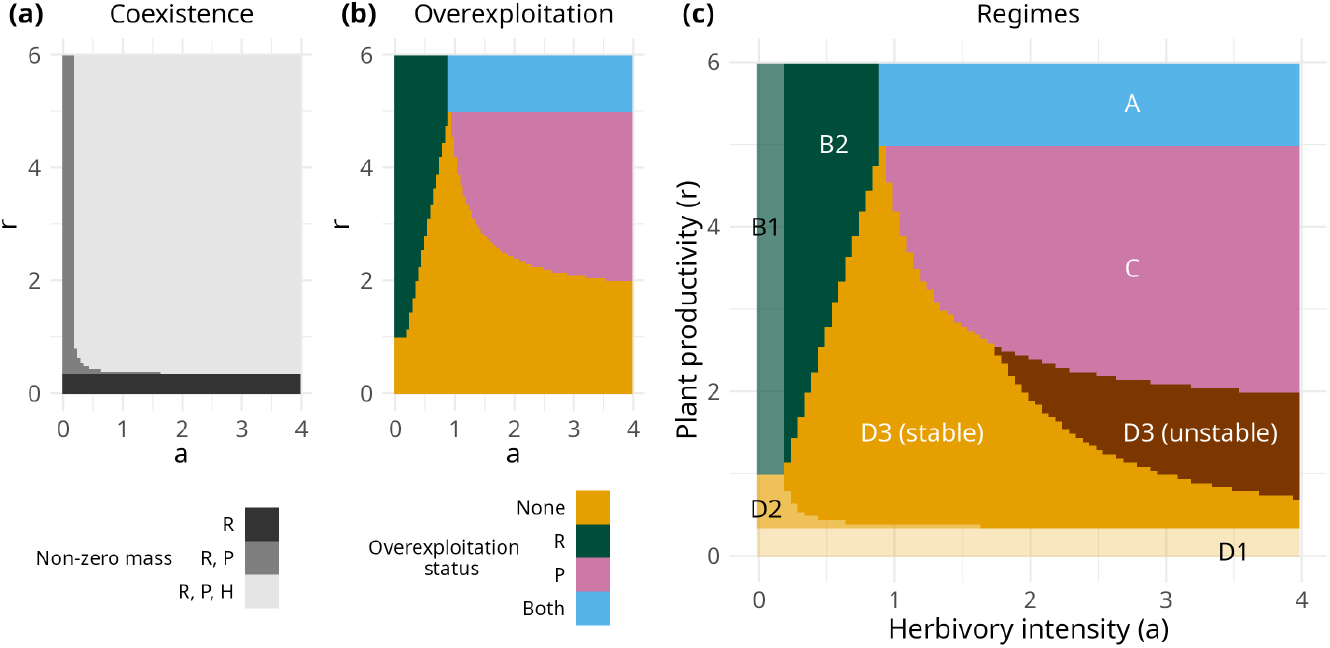
Trophic levels present at steady state (a), overexploitation status at steady state (b), and parameter domains of all possible regimes (c) while varying herbivory intensity, *a*, and plant productivity, *r*. The analytical expressions of nutrient masses at steady states of all three trophic levels for each attractor are given by equations S1 to S7. Letters in panel (c) indicate the analytical expression each domain corresponds to, with its stability in the case of the D3 regime. Parameter values other than a and r are found in Tab. 1.

### Biomass patterns with increasing interaction strength

Overexploitation of a given trophic level severely limits the top-down control exerted by its consumer. This general result appears directly in the equations of fixed points, where herbivore parameters explicitly set plant biomass, but only when plants are not overexploited (see Eqs. S1 and S3). Similarly, plant parameters only set resource biomass when resources are not overexploited (see Eqs. S3 and S4).

This general law translates into systematic variations in top-down and bottom-up controls, which we obtain directly from the signs of the derivatives of the different fixed points with respect to plant productivity and herbivory intensity (Tab. 2, see details in Tab. S1). As expected, classical top-down patterns are observed when no overexploitation occurs; that is, increasing plant productivity benefits the herbivore and not the top-down regulated plant, and increasing herbivory pressure produces anti-correlated changes of adjacent trophic levels (Tab. 2a). Such an outcome is conserved when the consumption of the overexploited level is not being manipulated (e.g. changing herbivory intensity while the resources are overexploited, Tab. 2b, or changing plant productivity while the plants are overexploited, Tab. 2c). Interestingly, when only plants are overexploited, raising plant productivity allows a coexistence of top-down and bottom-up controls along the food chain (Tab. 2c, olive color). Classical top-down variations are progressively eroded when overexploitation occurs, as highlighted by grey colors on Tab. 2.

**Table 2:**
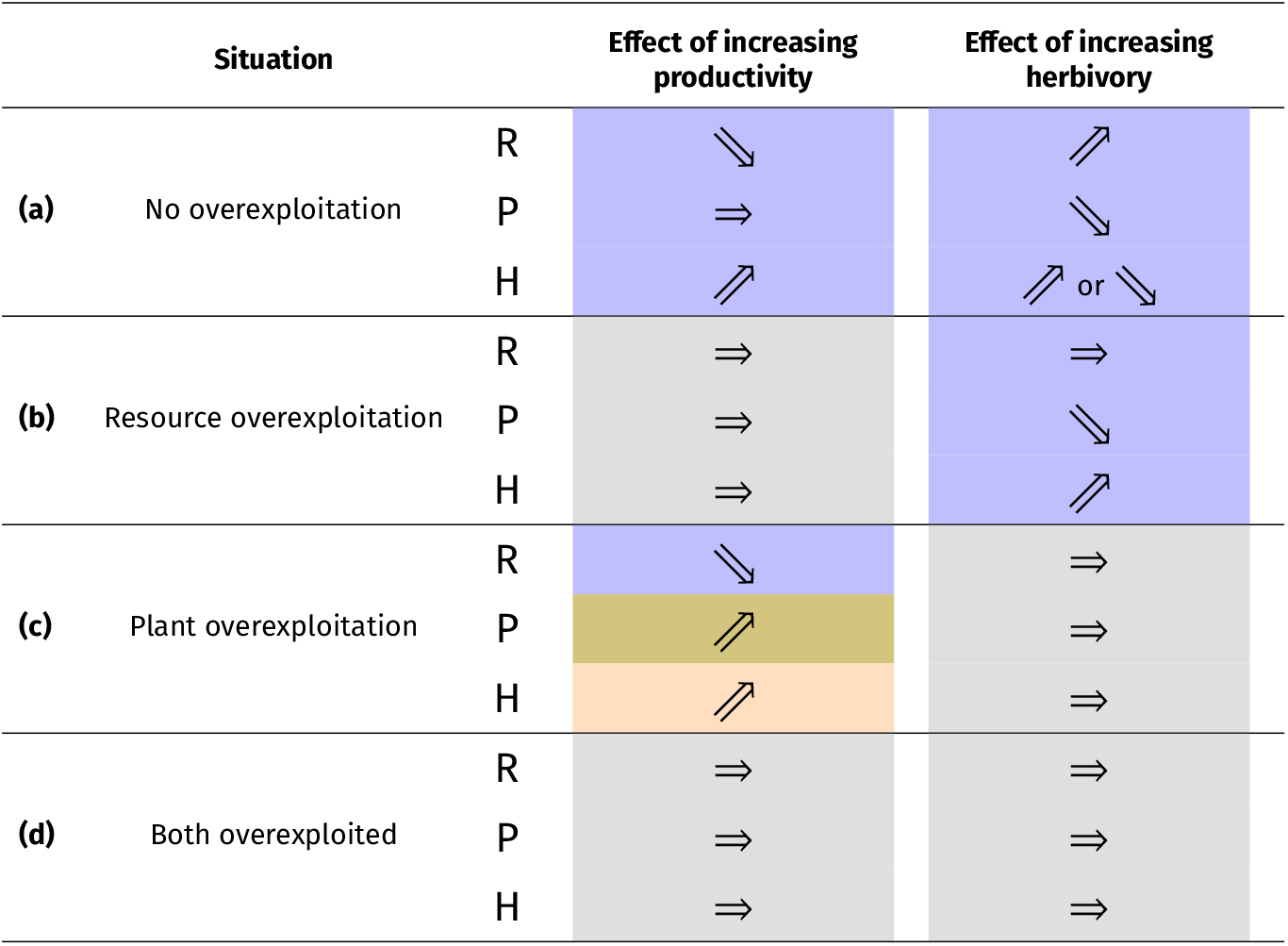
Biomass patterns caused by increasing plant productivity or herbivory intensity at steady states where the three trophic levels coexist. Colored cells denote biotic control type: top-down (blue), bottom-up with increasing biomass levels (yellow) and bottom-up with stagnating biomass levels (grey). The olive-colored cell denotes that, when plants are overexploited, they exert both bottom-up control on the herbivore and top-down control on the nutrient. Arrow directions are based on the signs of the partial derivatives of the fixed points’ analytical expressions (see Tab. S1 for details).

As intuitively expected, when increasing plant productivity, overexploitation of inorganic resources by plants introduces an upper limit to plant and herbivore biomasses, eventually suppressing top-down control of plants by herbivores (Tab. S1, Tab. 2b, Fig. 3). This suppression of a classical pattern in ecology (Oksanen and Oksanen, 2000) occurs both at high and at low herbivory intensity past a certain plant productivity (Fig. 3). For both herbivory levels, herbivore biomass first typically increases while plant biomass stays constant or near-constant (top-down control) and inorganic resource decreases until it becomes overexploited. Thereafter, top-down controls are eroded, and higher productivity no longer increases biomasses in the system. At high herbivory, this erosion happens after a parameter range where plants are overexploited and therefore exert both a bottom-up control on herbivores and and top-down control on resources.

**Figure 3:**
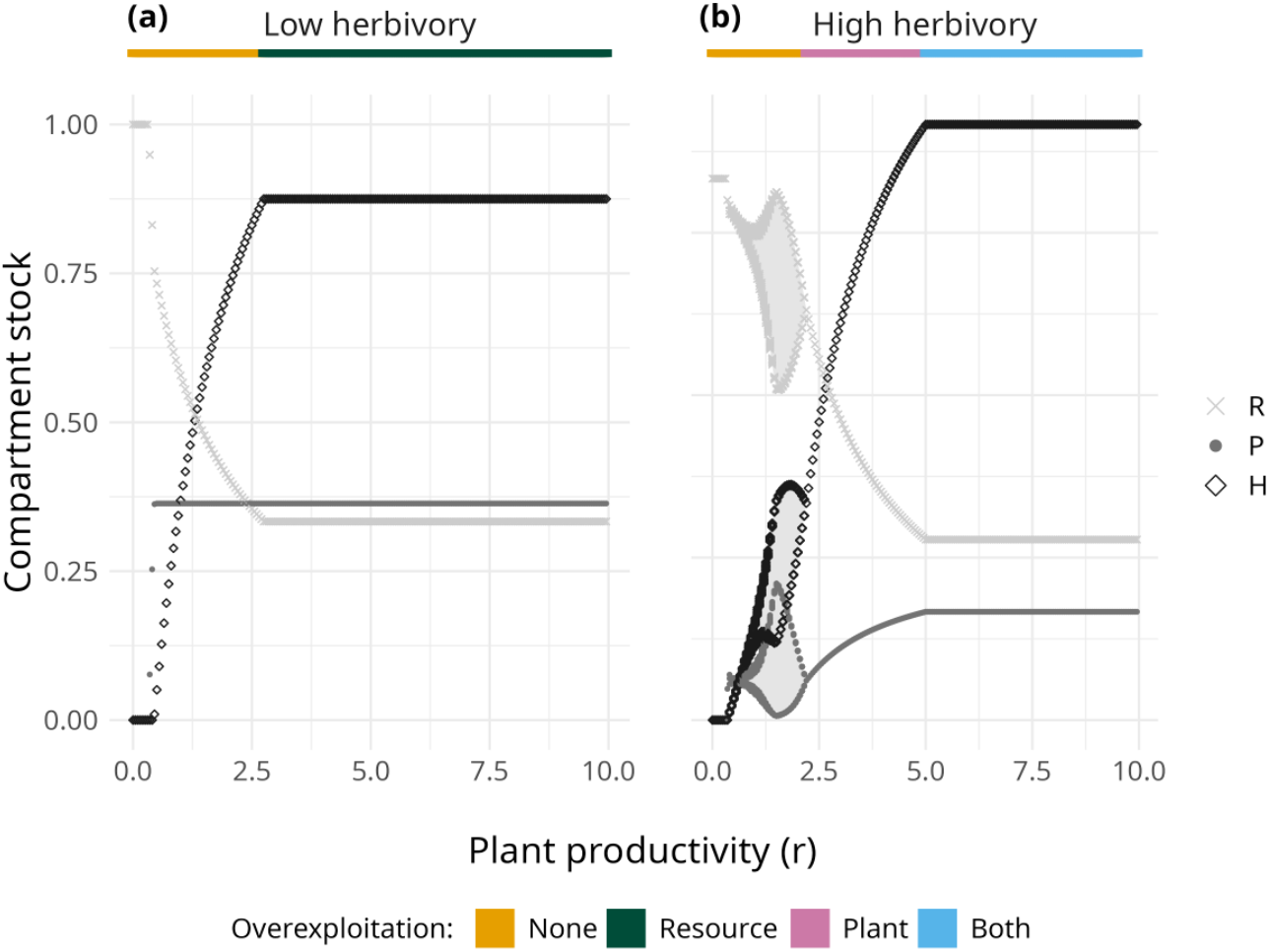
Equilibrium stocks of the three trophic levels along plant productivity for (a) low and (b) high herbivory intensities. The colored bar above the plot indicates the trophic levels under overexploitation. Shaded areas in panel (b) represent the ranges of each trophic level’s oscillations. *a* was set at 0.5 and 2.5 units for low and high herbivory, respectively. Other parameter values are found in Tab. 1.

Similarly, when increasing herbivory intensity, top-down control is eventually suppressed when the plant becomes overexploited (Tab. 2c, Fig. 4b and c), with herbivore biomass reaching a plateau at high herbivory. Note that when plant productivity is low, plant overexploitation does not occur because plant production cannot sustain the levels of herbivores that would lead to plant overexploitation (Fig. 4a). In these settings, a classical top-down pattern is first observed at low herbivory (correlated positive variations of herbivore and resource associated with a decrease in plant biomass around *a* = 0.2), but as herbivory increases the herbivore biomass starts to decline. This decline coincides with the herbivory level preventing the overexploitation of resources by plants, leading to such a decline of plants that it affects herbivore biomass. Intermediate values of plant productivity sustain higher herbivore biomass (Fig. 4b), that allows the overexploitation of plants by herbivores at high herbivory intensity, thus halting the herbivore decline observed on Fig. 4a. At high plant productivity, plants and herbivores reach sufficiently high levels to impose overexploitation of first resources, then both resources and plants with increasing herbivory, without the step where the herbivore biomass is sufficient to relax resource overexploitation via top-down control but too low to impose plant overexploitation (Fig. 4c).

**Figure 4:**
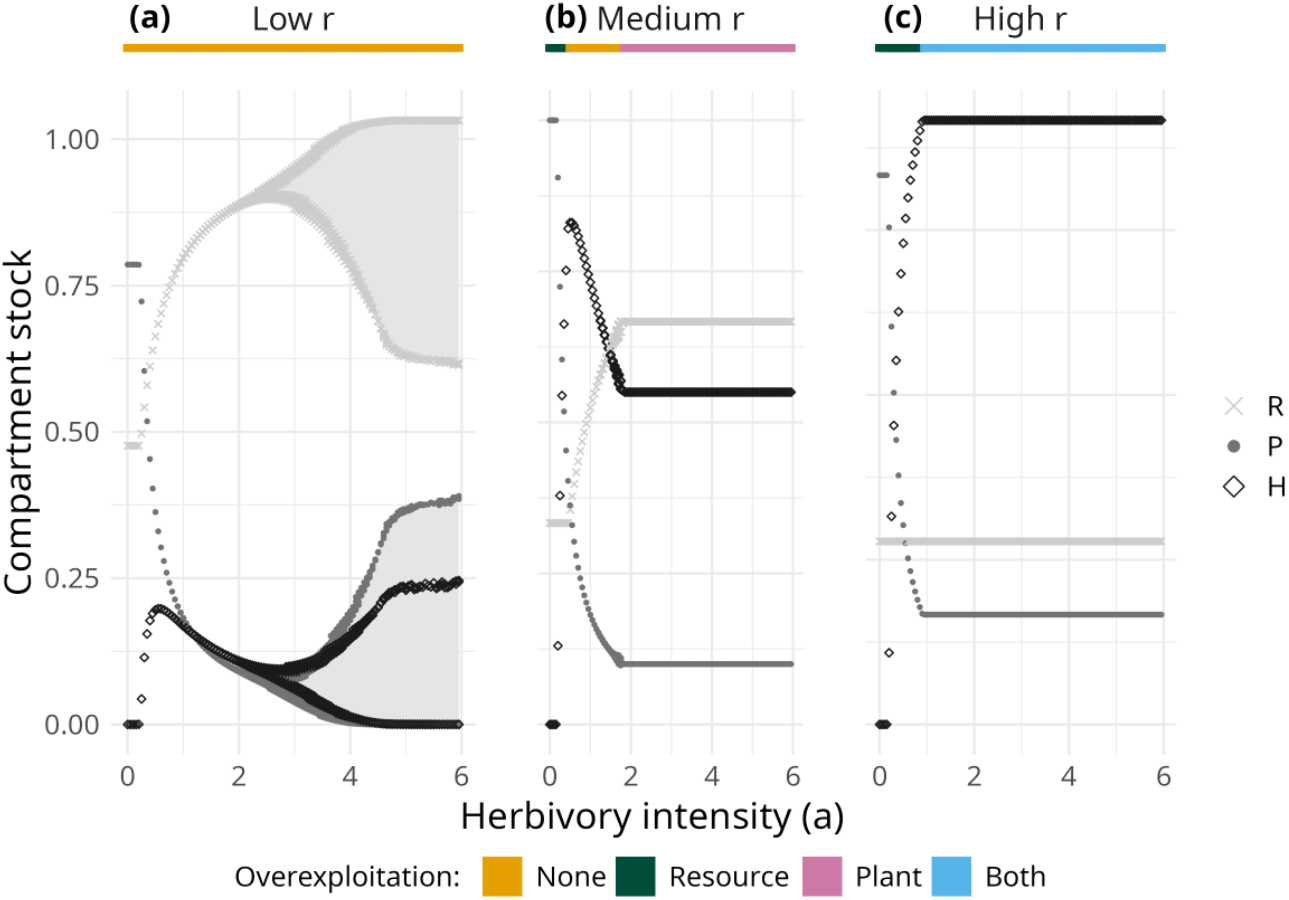
Equilibrium stocks of the three trophic levels along herbivory intensity for (a) low, (b) medium, and (c) high plant productivity. The colored bar above the plot indicates the trophic levels under overexploitation. Shaded areas in panel (a) represent the ranges of each trophic level’s oscillations. *r* was set at 0.7, 2.5 and 6 units for low, medium and high plant productivity, respectively. Other parameter values are found in Tab. 1.

### Stability, interaction strength and the paradox of enrichment

As expected, both the decrease in stability with interaction strength (Rip and McCann 2011) and the paradox of enrichment (Rosenzweig 1971) are often present without overexploitation but are suppressed by overexploitation as shown by the temporal dynamics of our model.

Without overexploitation, the stability of the system decreases both with plant productivity and with herbivory intensity, because increasing any of them increases energy fluxes, which can cause the appearance of oscillations. In this regime, no fluctuations happen at low productivity and herbivory (stable node). Increase in plant productivity or herbivory leads to damped oscillations (spiral node) and eventually to sustained cycles (brown on Fig. 2c). Particularly, increased plant productivity enhances energy fluxes, thereby creating an enrichment that is destabilizing (i.e. the paradox of enrichment). Such instabilities are suppressed when interaction strengths are so high that the plant and/or the resource become overexploited. For instance, the appearance of cycles at higher herbivory is observed in a situation where plant overexploitation does not occur (low plant productivity, Fig. 4a) but is suppressed at higher plant productivity, where sufficient herbivore stocks lead to plant overexploitation (Figs. 3b and 4b). When herbivory intensity is low, enrichment is not destabilizing (Fig. 3a).

## Discussion

While many instances of overexploitation in a variety of organisms and ecosystems have been documented, their implications for biomass variations along food chains are still largely unclear. Our work shows that while classical patterns expected from top-down controls apply without overexploitation, and may still happen in particular overexploitation situations, these patterns are most often suppressed, especially when overexploitation happens for both resources and plants. Moreover, the destabilization classically found when increasing productivity or herbivory is dampened by overexploitation: under overexploitation, further increasing them does not lead to oscillating dynamics. We discuss how these results relate to existing literature on biotic control and ecosystem stability.

The idea of that the growth of an organism requires a minimum amount of resources has been a long-standing assumption in studies focusing on nutrient availability to phytoplankton cell cultures, since the formulation of Droop’s variable internal stores model (Droop 1974). In this framework, cell growth ceases if the internal concentration of a nutrient falls below a certain threshold, and algae growth can be limited by several nutrients, ultimately affecting population growth. This framework has been applied in studies of competition between algae (Tilman 1977), where the presence of a competitor lowers the available nutrient quantity and therefore decreases the population of an algae of interest, stressing the importance of bottom-up control in the shaping of algal communities. In this study, we expand the idea of a threshold that can limit consumer growth to a more generic context where overexploitation is applied at the population rather than at the individual scale and can occur at different trophic levels.

### Over-exploitation suppresses top-down control

Accounting for overexploitation explicitly, we show some unexpected relationships between the masses at equilibrium for the different compartments and the strengths of trophic interactions. Given classical top-down controls in food chains, herbivore biomass only stops increasing with primary productivity when productivity allows the existence of a predator level (Oksanen and Oksanen 2000), just like plant biomass is capped when productivity allows the existence of a herbivore (without predators). Our results show that, without carnivores, herbivore biomass can be capped when plants start overexploiting their resources (Fig. 3a), shifting the top-down to a bottom-up regulation of herbivores. Indeed, even if they are highly productive (i.e. with a high potential resource consumption), the plants harvest all available resource mass at every time step. The plants are therefore under bottom-up control and their biomass production cannot exceed a certain level, which in turn sets an upper boundary for herbivore biomass. Similarly, herbivory pressure is classically expected to limit the plant and to allow for a larger inorganic resource pool (i.e. a trophic cascade). Again, in our model, this effect is limited by plant overexploitation. Fig. 4b for instance shows the expected decrease in plant biomass and increases in inorganic resource stocks and herbivore biomass when herbivory intensity increases, but only up to the point where herbivores overexploit the plants, suppressing the expected top-down pattern.

Empirically, bottom-up controlled food chains have mostly been reported in terrestrial or nutrient-limited marine ecosystems while top-down controlled chains are reported to dominate in freshwater ecosystems (Shurin et al. 2002). For example, in a manipulative experiment on the effect of light pollution on terrestrial plant communities, (Anic et al. 2022) increasing the photoperiod of nutrient-limited plant communities (therefore increasing their possible productivity) does not lead to significant plant biomass increase, suggesting a clear bottom-up control of plants by resource availability. Conversely, (Su et al. 2021) report a majority of typical top-down trophic cascades in a meta-analysis of fish addition experiments to freshwater communities. The time scale of regeneration has already been suggested as a major driver of the difference in biotic control between freshwater and terrestrial ecosystems (Chase 2000), with for example faster decomposition rates in freshwater systems (Gounand et al. 2020). Our study contributes to this argument by showing that, when resource regeneration is not sufficient (i.e. when there is overexploitation), bottom-up control dominates the food chain.

Our model shows a simple way of introducing bottom-up control in certain situations for a system that otherwise behaves as top-down controlled. The use of a food chain model with three distinct trophic levels and only one directed resource flow between each pair of trophic levels does not allow us to incorporate much heterogeneity in trophic control (i.e. bottom-up and top-down control co-occuring in a food web, following Hunter and Price 1992). Considering different functional groups with different size classes and diets has been shown to confer upon the food web more biotic control heterogeneity (Hulot et al. 2000). We anticipate that incorporating different overexploitation requirements between functional groups of the same trophic level in our model, such as those induced by differences in resource accessibility or preferability will provide new insights on coexistence and biomass variations within food webs. Interestingly, our results show that overexploitation still allows a situation where plants exert both top-down and bottom-up controls in a simple food chain: when it is overexploited but still under increasing productivity (Fig. 3b, Tab. 2). While we do not consider heterogeneity of control within trophic levels, we can still observe it at the scale of the whole food chain.

More recently, the scaling of interaction strength along the food chain has been proposed as an important determinant for the type of biotic control (Barbier and Loreau 2019, Galiana et al. 2021). In these studies, top-down control is associated with situations where the interaction strength scales with consumer metabolism, while in bottom-up control interaction strength scales with resource metabolism. Our model implements a similar way of exploring bottom-up and top-down controls in food chains: overexploitation relates to situations where the interaction is limited by the regeneration of the resource (i.e. its metabolism) and we indeed show that overex-ploitation is mostly associated with bottom-up controls. Directly comparing the two frameworks would require to convert the nutrient masses in our model to biomass stocks (and thus to biomass pyramids as in Barbier and Loreau 2019), which would rely on assumptions for the ratio between nutrient mass and biomass at each trophic level.

### Overexploitation counteracts the paradox of enrichment

Conditions of stability of trophic systems have received a lot of attention (Rosenzweig 1971, Pimm and Lawton 1977, McCann et al. 1998, Rip and McCann 2011), as less stable systems may often lead to extinctions of species due to large fluctuations in densities, or to a low resilience to external disturbances. Particularly, while high energy flows are usually supposed to help satisfying the needs of each species and to help maintaining them, they may also lead to unstable (and therefore fragile) systems (Rosenzweig 1971, Rip and McCann 2011). Under overexploitation (facilitating bottom-up control), we do not observe this paradox of enrichment while our model reproduces it when there is no overexploitation (i.e. under top-down control). A change in species productivity or consumption pressure that switches our system from no overexploitation to overexploitation can thus suppress the paradox of enrichment in a resource-consumer system where it was previously observed. This may contribute to the debate around the discrepancy between enrichment theory and empirical evidence (Jensen and Ginzburg 2005). Indeed, as reviewed by Roy and Chattopadhyay 2007, empirical studies have both documented situations where the paradox of enrichment is observed and situations where it is not. For example, in rather similar zooplankton-algae experimental systems, Fussmann et al. 2000 and Kirk 1998 respectively found evidence for destabilization and for stabilization by enrichment. Ratio-dependent population models (Arditi and Ginzburg 1989), where the functional response depends on the ratio between resource and consumer densities, while they suppress the paradox of enrichment, do not reproduce it in any condition. Interestingly, our model reproduces situations where the paradox of enrichment is suppressed while still using prey-dependent functional responses, as widely used in population dynamics models. Moreover, since overexploitation occurs when there is a difference in time scales between resource dynamics and consumer feeding, our model generates the testable prediction that the paradox of enrichment should be less likely to occur when the regeneration of a prey is slower than the feeding ability of its consumer.

Similarly, Rip and McCann 2011 report a destabilizing effect of the strength of trophic interactions, but they do so in the top-down controlled Lotka-Volterra and Rosenzweig models. Over-exploitation in our model then alleviates this effect of the consumer on its resource and shows situations where further increasing interaction strength does not decrease stability (Fig. 4b and c). Overall, our results suggest that overex-ploitation might contribute to the temporal stability of ecosystems.

### Further investigations

While our model explicitly includes recycling, we only explore the possibility of reduced or suppressed recycling and its impact on the behavior of our model as a robustness analysis (Fig. S3), in which we checked that the overexploitation situations did not qualitatively change with recycling strength. However, our implementation of recycling makes it constant along the food chain, without introducing a different scaling between herbivore biomass and plant biomass recycling, which would likely have an impact on system properties (Mazancourt et al. 1998). The role of the scaling between recycling and interaction terms on observed patterns could directly be investigated, possibly altering the prevalence of the different regimes. We predict that highly efficient recycling would hinder resource overexploitation, but the way this quantitatively influences the steady state of the system remains to be tested.

The interaction of herbivory, recycling and overexploitation patterns could also lead to the emergence of spatial patterns at all trophic levels, leading to meta-ecosystems (Loreau et al. 2003). Source and sink dynamics and nutrient exports reported in meta-ecosystems (Gravel et al. 2010, Loreau et al. 2013, Gounand et al. 2014) should have a strong impact in a spatialized equivalent of our model, because dispersing herbivores or plants may be able to escape local overexploitation of their resources. By modulating food availability to the herbivore, spatial processes and their metabolic cost may alter the balance between energy dissipation and acquisition, possibly changing the way energy is transferred along the food chain. Acquiring a proper understanding of the local system here was however crucial before bringing the study of overexploitation to a spatially explicit context.

Finally, we only focused on the ecological implications of overexploitation, leaving out the possibility of eco-evolutionary dynamics. Overexploitation however corresponds to strong selective pressures within trophic levels. For instance, plants facing resource overexploitation would be expected to evolve in resource acquisition or competitive traits. Overexploitation of plants by herbivores may yield strong selection on their foraging traits or behaviors and on the defensive phenotypes of plants. Interestingly, the evolution of defenses has been shown to decrease top-down controls in food chains (Loeuille and Loreau 2004), yielding a dominance of bottom-up patterns. In turn, empirical works suggest that such defended or unpalatable plants largely reduce the possibility of overexploitation by their consumers (White 2005).

We here suggest that overexploitation can affect our qualitative view of trophic control and stability in natural ecosystems. Because high nutrient flows (McCann et al. 2021), changes in plant productivity (O’Sullivan et al. 2020), and overgrazing (Bertness et al. 2014, Vergés et al. 2014, Archer et al. 2017, Maestre et al. 2022) are all important current changes, we hope that our study will stimulate empirical and experimental tests of the effects of overexploitation to better anticipate future possible scenarios.

## Significance statement

Overexploitation of plants or nutrients is a major component of the functioning of many natural ecosystems, but is often overlooked in consumer-resource theory. Using a simple model, we show that when overexploitation occurs at high interaction strength it can suppress well-known but debated phenomena in ecology: the prevalence of top-down control and the destabilization of population dynamics by high productivity leading to the paradox of enrichment. These are observed when consumption pace still allows the resource to regenerate, but are suppressed under overexploitation. These findings connect contrasting views about trophic interactions in ecology and contribute to the reconciliation between long-standing divergent paradigms.

## Code availability

All Julia and R code required to run simulations and generate figures with their corresponding documentation are currently private for peer-review. However, they can already be requested from the authors via email and they will be public after peerreview, as a permanent archive at https://doi.org/10.5061/dryad.ffbg79d4g and as a public GitHub repository.

## Author contributions

Conceptualization and Methodology: JG, NL and IG. Formal Analysis, Software and Visualization: JG. Writing - original draft: JG. Writing - review & editing: JG, NL and IG.

## Supporting information for

**Figure S1:**
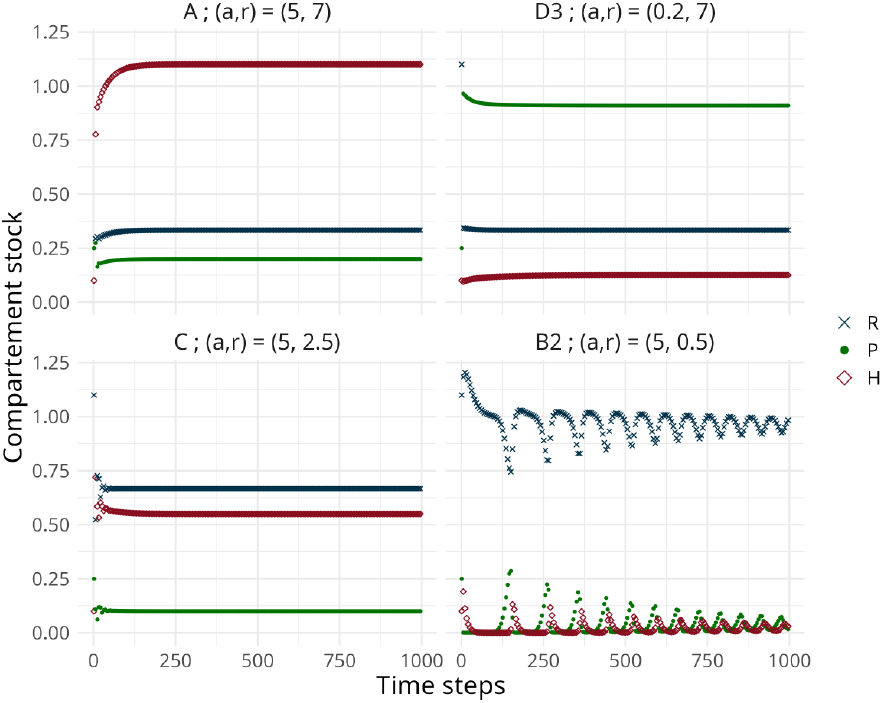
Examples of time series in regimes with herbivores. Herbivory intensity and plant productivity parameters used to generate the time series are shown on panel labels. Other parameter values are found in Tab. 1.

**Figure S2:**
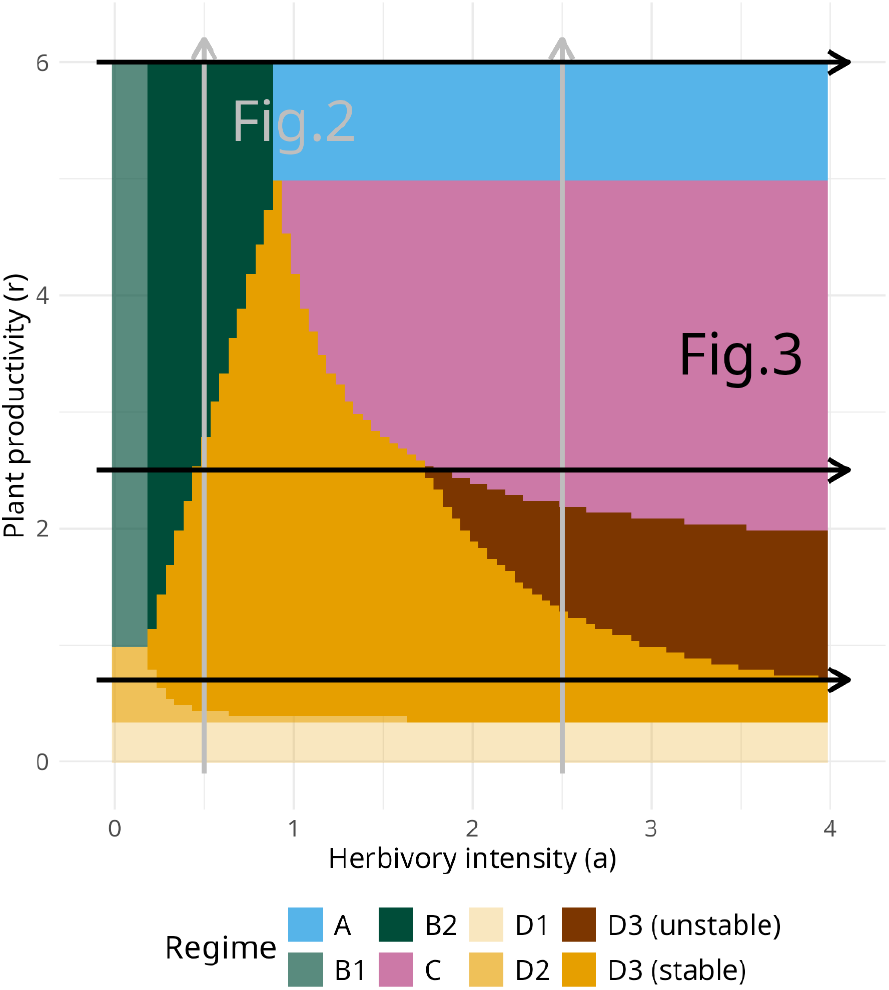
Parameter ranges in Figs. 3 and 4 (arrows) as transects across the parameter space. Studied transects offer a good overview of the different possible transitions between regimes and overex-ploitation status. See Results for a detailed description of biomass patterns along transects.

**Figure S3:**
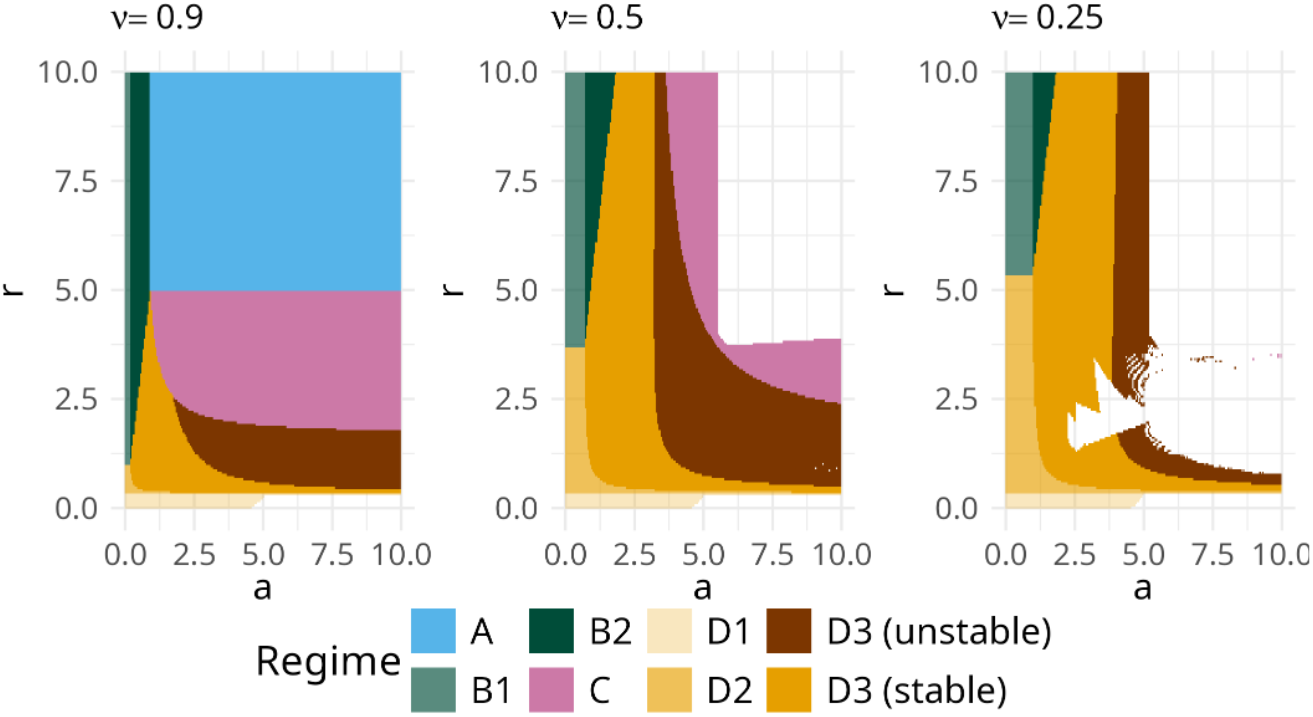
Parameter domains for possible regimes for decreased recycling efficiency ν. Blank spaces correspond to parameter ranges where the simulations reach negative values. Further decreasing ν after 0.5 leads to more blank areas but still yields similar domains and bifurcations.

### Analytical expressions for the fixed points

Analytical expressions for the eight fixed points are given by Equations S1 to S7, with the fixed point’s name in indices of the state variables (A, B1, B2, C, D1, D2, D3). In each system of equations, overexploited compartments are shown with underlined symbols. Overexploitation of both plant and resource leads to a single stable fixed point (Eq. S1):

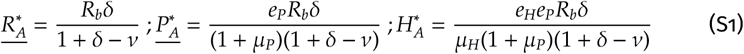

Overexploitation of the resource alone led to the exclusion of herbivores (Eq. S2), or to a stable fixed point with herbivores when *a* was high enough (Eq. S3):

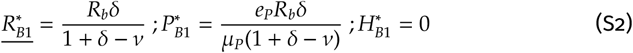

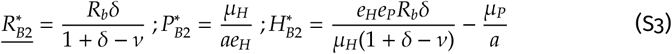

Forcing overexploitation of the plant alone led to a stable fixed point with both plant and herbivore (Eq. S4) and a fixed point without herbivores nor plants, which is never valid because it does not satisfy the validity condition for plant overexploitation.

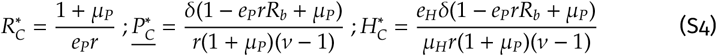

Without any overexploitation, there were 3 fixed points. One excluded both plant and herbivores (Eq. S5), one excluded the herbivores, (Eq. S6), and a third fixed point showed coexistence of resource, plant and herbivore (Eq. S7).

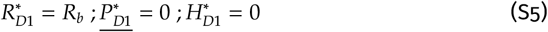

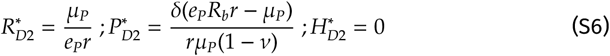

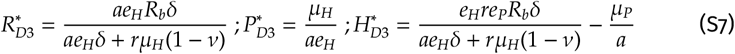

### Effect of increasing *r* **and** *a*

**Table S1:**
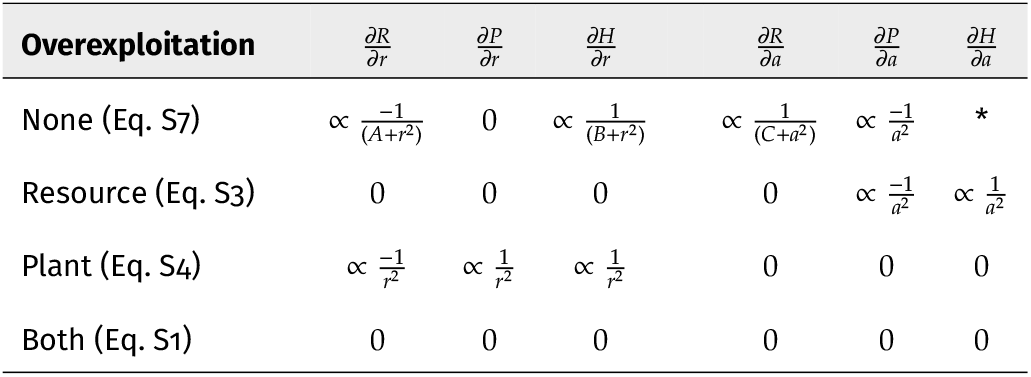
Signs of the derivatives of steady-state biomass expressions with respect to *r* and to *a*. Only steady states with coexisting plants and herbivores are investigated. *A, B* and *C* are positive constants that depend on other model parameters. The derivative (*) at the top left of the table does not have a constant sign for all *a*. In the parameter ranges used in our simulations, it is positive when *a* is small and negative when *a* is high (Fig. 4).

| Overexploitation | $\frac{\partial R}{\partial r}$ | $\frac{\partial P}{\partial r}$ | $\frac{\partial H}{\partial r}$ | $\frac{\partial R}{\partial a}$ | $\frac{\partial P}{\partial a}$ | $\frac{\partial H}{\partial a}$ |
| --- | --- | --- | --- | --- | --- | --- |
| None (Eq. S7) | $\propto \frac{-1}{(A+r^2)}$ | 0 | $\propto \frac{1}{(B+r^2)}$ | $\propto \frac{1}{(C+a^2)}$ | $\propto \frac{-1}{a^2}$ | * |
| Resource (Eq. S3) | 0 | 0 | 0 | 0 | $\propto \frac{-1}{a^2}$ | $\propto \frac{1}{a^2}$ |
| Plant (Eq. S4) | $\propto \frac{-1}{r^2}$ | $\propto \frac{1}{r^2}$ | $\propto \frac{1}{r^2}$ | 0 | 0 | 0 |
| Both (Eq. S1) | 0 | 0 | 0 | 0 | 0 | 0 |

## References

Anic, Vinka, Kevin J. Gaston, Thomas W. Davies, and Jonathan Bennie. 2022. “Long-Term Effects of Artificial Nighttime Lighting and Trophic Complexity on Plant Biomass and Foliar Carbon and Nitrogen in a Grassland Community”. Ecol. Evol. 12.8, e9157. doi:10.1002/ece3.9157.

Archer, Steven R, Erik M Andersen, Katharine I Predick, Susanne Schwinning, Robert J. Steidl, and Steven R Woods. 2017. “Woody Plant Encroachment: Causes and Consequences”. Rangeland Systems: Processes, Management and Challenges. Ed. by David D. Briske, pp. 25–84.

Arditi, Roger and Lev R. Ginzburg. 1989. “Coupling in Predator-Prey Dynamics: Ratio-Dependence”. J. Theor. Biol. 139.3, pp. 311–326. doi:10.1016/S0022-5193(89)80211-5.

Barbier, Matthieu and Michel Loreau. 2019. “Pyramids and Cascades: A Synthesis of Food Chain Functioning and Stability”. Ecol. Lett.22.2. Ed. by Hillary Young, pp. 405–419. doi:10.1111/ele.13196.

Bertness, Mark D., Caitlin P. Brisson, Tyler C. Coverdale, Matt C. Bevil, Sinead M. Crotty, and Elena R. Suglia. 2014. “Experimental Predator Removal Causes Rapid Salt Marsh Die-Off”. Ecol. Lett. 17.7, pp. 830–835. doi:10.1111/ele.12287.

Bezanson, Jeff, Stefan Karpinski, Viral B. Shah, and Alan Edelman. 2012. Julia: A Fast Dynamic Language for Technical Computing. doi:10.48550/ARXIV.1209.5145. 1209.5145 [cs].

Bideault, Azenor, Núria Galiana, Yuval R. Zelnik, Dominique Gravel, Michel Loreau, Matthieu Barbier, and Arnaud Sentis. 2021. “Thermal Mismatches in Biological Rates Determine Trophic Control and Biomass Distribution under Warming”. Glob. Change Biol. 27.2, pp. 257–269. doi:10.1111/gcb.15395.

Browning, Thomas J. and C. Mark Moore. 2023. “Global Analysis of Ocean Phytoplankton Nutrient Limitation Reveals High Prevalence of Co-Limitation”. Nat. Commun. 14.1, p. 5014. doi:10.1038/s41467-023-40774-0.

Chase, Jonathan M. 2000. “Are There Real Differences among Aquatic and Terrestrial Food Webs?” Trends Ecol. Evol. 15.10, pp. 408–412. doi:10.1016/S0169-5347(00)01942-X.

Christianen, Marjolijn J. A., Peter M. J. Herman, Tjeerd J. Bouma, Leon P. M. Lamers, Marieke M. van Katwijk, Tjisse van der Heide, Peter J. Mumby, Brian R. Silliman, Sarah L. Engelhard, Madelon van de Kerk, Wawan Kiswara, and Johan van de Koppel. 2014. “Habitat Collapse Due to Overgrazing Threatens Turtle Conservation in Marine Protected Areas”. Proc. R. Soc. B 281.1777, p. 20132890. doi:10.1098/rspb.2013.2890.

Datseris, George. 2018. “DynamicalSystems.Jl: A Julia Software Library for Chaos and Nonlinear Dynamics”. J. Open Source Softw. 3.23, p. 598. doi:10.21105/joss.00598.

Droop, Michael R. 1968. “Vitamin B12 and Marine Ecology. IV. The Kinetics of Uptake, Growth and Inhibition in Monochrysis Lutheri”. J. Mar. Biol. Assoc. U. K. 48.3, pp. 689–733. doi:10.1017/S0025315400019238.

Droop, Michael R. 1974. “The Nutrient Status of Algal Cells in Continuous Culture”. J. Mar. Biol. Assoc. U. K. doi:10.1017/S002531540005760X.

Elser, James J., Matthew E.S. Bracken, Elsa E. Cleland, Daniel S. Gruner, W. Stanley Harpole, Helmut Hillebrand, Jacqueline T. Ngai, Eric W. Seabloom, Jonathan B. Shurin, and Jennifer E. Smith. 2007. “Global Analysis of Nitrogen and Phosphorus Limitation of Primary Producers in Freshwater, Marine and Terrestrial Ecosystems”. Ecol. Lett. 10.12, pp. 1135–1142. doi:10.1111/j.1461-0248.2007.01113.x.

Elton, Charles S. 1927. Animal Ecology. 2nd edition, 2001. University of Chicago Press.

Forbes, Elizabeth S., J. Hall Cushman, Deron E. Burkepile, Truman P. Young, Maggie Klope, and Hillary S. Young. 2019. “Synthesizing the Effects of Large, Wild Herbivore Exclusion on Ecosystem Function”. Funct. Ecol. 33.9, pp. 1597–1610. doi:10.1111/1365-2435.13376.

Frederiksen, Morten, Martin Edwards, Anthony J. Richardson, Nicholas C. Halliday, and Sarah Wanless. 2006. “From Plankton to Top Predators: Bottom-up Control of a Marine Food Web across Four Trophic Levels”. J. Anim. Ecol. 75.6, pp. 1259–1268. doi:10.1111/j.1365-2656.2006.01148.x.

Fussmann, Gregor F., Stephen P. Ellner, Kyle W. Shertzer, and Nelson G. Hairston Jr. 2000. “Crossing the Hopf Bifurcation in a Live Predator-Prey System”. Science 290.5495, pp. 1358–1360. doi:10.1126/science.290.5495.1358.

Galiana, Núria, Jean-François Arnoldi, Matthieu Barbier, Amandine Acloque, Claire de Mazancourt, and Michel Loreau. 2021. “Can Biomass Distribution across Trophic Levels Predict Trophic Cascades?” Ecol. Lett. 24.3, pp. 464–476. doi:10.1111/ele.13658.

Gounand, Isabelle, Chelsea J. Little, Eric Harvey, and Florian Altermatt. 2020. “Global Quantitative Synthesis of Ecosystem Functioning across Climatic Zones and Ecosystem Types”. Global Ecol. Biogeogr. 29.7, pp. 1139–1176. doi:10.1111/geb.13093.

Gounand, Isabelle, Nicolas Mouquet, Elsa Canard, Frédéric Guichard, Céline Hauzy, and Dominique Gravel. 2014. “The Paradox of Enrichment in Metaecosystems”. Am. Nat. 184.6, pp. 752–763. doi:10.1086/678406.

Gravel, Dominique, Frédéric Guichard, Michel Loreau, and Nicolas Mouquet. 2010. “Source and Sink Dynamics in Meta-Ecosystems”. Ecology 91.7, pp. 2172–2184. doi:10.1890/09-0843.1.

Hulot, Florence D., Gérard Lacroix, Françoise Lescher-Moutoué, and Michel Loreau. 2000. “Functional Diversity Governs Ecosystem Response to Nutrient Enrichment”. Nature 405.6784, pp. 340–344.doi: 10.1038/35012591.

Hunter, Mark D. and Peter W. Price. 1992. “Playing Chutes and Ladders: Heterogeneity and the Relative Roles of Bottom-Up and Top-Down Forces in Natural Communities”. Ecology 73.3, pp. 724–732. JSTOR: 1940152.

Jensen, Christopher X. J. and Lev R. Ginzburg. 2005. “Paradoxes or Theoretical Failures? The Jury Is Still Out”. Ecol. Modell. 188.1, pp. 3–14. doi:10.1016/j.ecolmodel.2005.05.001.

Kallio, Paavo and Juhani Lehtonen. 1975. “On the Ecocatastrophe of Birch Forests Caused by Oporinia Autumnata (Bkh.) and the Problem of Reforestation”. Fennoscandian Tundra Ecosystems: Part 2 Animals and Systems Analysis. Ed. by F. E. Wielgolaski. Berlin, Heidelberg: Springer, pp. 174–180.doi: 10.1007/978-3-642-66276-8_23.

Kamata, Naoto. 2002. “Outbreaks of Forest Defoliating Insects in Japan, 1950–2000”. Bull. Entomol. Res. 92.2, pp. 109–117. doi:10.1079/BER2002159.

Kirk, Kevin L. 1998. “Enrichment Can Stabilize Population Dynamics: Autotoxins and Density Dependence”. Ecology 79.7, pp. 2456–2462. doi:10.1890/0012-9658(1998)079[2456:ECSPDA]2.0.CO;2.

Lampert, Winfried, Walter Fleckner, Hakumat Rai, and Barbara E. Taylor. 1986. “Phytoplankton Control by Grazing Zooplankton: A Study on the Spring Clear-Water Phase1”. Limnol. Oceanogr. 31.3, pp. 478–490. doi:10.4319/lo.1986.31.3.0478.

Lindeman, Raymond L. 1942. “The Trophic-Dynamic Aspect of Ecology”. Ecology 23.4, pp. 399–417. doi: 10.2307/1930126.

Loeuille, Nicolas and Michel Loreau. 2004. “Nutrient Enrichment and Food Chains: Can Evolution Buffer Top-down Control?” Theor. Popul Biol. 65.3, pp. 285–298. doi:10.1016/j.tpb.2003.12.004.

Loreau, Michel, Tanguy Daufresne, Andrew Gonzalez, Dominique Gravel, Frédéric Guichard, Shawn J. Leroux, Nicolas Loeuille, François Massol, and Nicolas Mouquet. 2013. “Unifying Sources and Sinks in Ecology and Earth Sciences”. Biol. Rev. 88.2, pp. 365–379. doi:10.1111/brv.12003.

Loreau, Michel, Nicolas Mouquet, and Robert D. Holt. 2003. “Meta-Ecosystems: A Theoretical Framework for a Spatial Ecosystem Ecology”. Ecol. Lett. 6.8, pp. 673–679. doi:10.1046/j.1461-0248.2003.00483.x.

Lotka, Alfred James. 1925. Elements of Physical Biology. Williams & Wilkins.

Maestre, Fernando T., Yoann Le Bagousse-Pinguet, Manuel Delgado-Baquerizo, David J. Eldridge, Hugo Saiz, Miguel Berdugo, Beatriz Gozalo, Victoria Ochoa, Emilio Guirado, Miguel García-Gómez, Enrique Valencia, Juan J. Gaitán, Sergio Asensio, Betty J. Mendoza, César Plaza, Paloma Díaz-Martínez, Ana Rey, Hang-Wei Hu, Ji-Zheng He, Jun-Tao Wang, Anika Lehmann, Matthias C. Rillig, Simone Cesarz, Nico Eisenhauer, Jaime Martínez-Valderrama, Eduardo Moreno-Jiménez, Osvaldo Sala, Mehdi Abedi, Negar Ahmadian, Concepción L. Alados, Valeria Aramayo, Fateh Amghar, Tulio Arredondo, Rodrigo J. Ahumada, Khadijeh Bahalkeh, Farah Ben Salem, Niels Blaum, Bazartseren Boldgiv, Matthew A. Bowker, Donaldo Bran, Chongfeng Bu, Rafaella Canessa, Andrea P. Castillo-Monroy, Helena Castro, Ignacio Castro, Patricio Castro-Quezada, Roukaya Chibani, Abel A. Conceição, Courtney M. Currier, Anthony Darrouzet-Nardi, Balázs Deák, David A. Donoso, Andrew J. Dougill, Jorge Durán, Batdelger Erdenetsetseg, Carlos I. Espinosa, Alex Fajardo, Mohammad Farzam, Daniela Ferrante, Anke S. K. Frank, Lauchlan H. Fraser, Laureano A. Gherardi, Aaron C. Greenville, Carlos A. Guerra, Elizabeth Gusmán-Montalvan, Rosa M. Hernández-Hernández, Norbert Hölzel, Elisabeth Huber-Sannwald, Frederic M. Hughes, Oswaldo Jadán-Maza, Florian Jeltsch, Anke Jentsch, Kudzai F. Kaseke, Melanie Köbel, Jessica E. Koopman, Cintia V. Leder, Anja Linstädter, Peter C. le Roux, Xinkai Li, Pierre Liancourt, Jushan Liu, Michelle A. Louw, Gillian Maggs-Kölling, Thulani P. Makhalanyane, Oumarou Malam Issa, Antonio J. Manzaneda, Eugene Marais, Juan P. Mora, Gerardo Moreno, Seth M. Munson, Alice Nunes, Gabriel Oliva, Gastón R. Oñatibia, Guadalupe Peter, Marco O. D. Pivari, Yolanda Pueyo, R. Emiliano Quiroga, Soroor Rahmanian, Sasha C. Reed, Pedro J. Rey, Benoit Richard, Alexandra Rodríguez, Víctor Rolo, Juan G. Rubalcaba, Jan C. Ruppert, Ayman Salah, Max A. Schuchardt, Sedona Spann, Ilan Stavi, Colton R. A. Stephens, Anthony M. Swemmer, Alberto L. Teixido, Andrew D. Thomas, Heather L. Throop, Katja Tielbörger, Samantha Travers, James Val, Orsolya Valkó, Liesbeth van den Brink, Sergio Velasco Ayuso, Frederike Velbert, Wanyoike Wamiti, Deli Wang, Lixin Wang, Glenda M. Wardle, Laura Yahdjian, Eli Zaady, Yuanming Zhang, Xiaobing Zhou, Brajesh K. Singh, and Nicolas Gross. 2022. “Grazing and Ecosystem Service Delivery in Global Drylands”. Science 378.6622, pp. 915–920. doi:10.1126/science.abq4062.

Mazancourt, Claire De, Michel Loreau, and Luc Abbadie. 1998. “GRAZING OPTIMIZATION AND NUTRIENT CYCLING: WHEN DO HERBIVORES ENHANCE PLANT PRODUCTION?” Ecology 79.7, p. 11. doi:10.1890/0012-9658(1998)079[2242:GOANCW]2.0.CO;2.

McCann, Kevin, Kevin Cazelles, Andrew S. MacDougall, Gregor F. Fussmann, Carling Bieg, Melania Cristescu, John M. Fryxell, Gabriel Gellner, Brian Lapointe, and Andrew Gonzalez. 2021. “Landscape Modification and Nutrient-Driven Instability at a Distance”. Ecol. Lett. 24.3, pp. 398–414. doi:10.1111/ele.13644.

McCann, Kevin, Alan Hastings, and Gary R. Huxel. 1998. “Weak Trophic Interactions and the Balance of Nature”. Nature 395.6704, pp. 794–798. doi:10.1038/27427.

McCauley, Douglas J., Gabriel Gellner, Neo D. Martinez, Richard J. Williams, Stuart A. Sandin, Fiorenza Micheli, Peter J. Mumby, and Kevin S. McCann. 2018. “On the Prevalence and Dynamics of Inverted Trophic Pyramids and Otherwise Top-Heavy Communities”. Ecol. Lett. 21.3, pp. 439–454. doi:10.1111/ele.12900.

Murray, James D. 2002. Mathematical Biology. 3rd ed. New York: Springer.

O’Sullivan, Michael, William K. Smith, Stephen Sitch, Pierre Friedlingstein, Vivek K. Arora, Vanessa Haverd, Atul K. Jain, Etsushi Kato, Markus Kautz, Danica Lombardozzi, Julia E. M. S. Nabel, Hanqin Tian, Nicolas Vuichard, Andy Wiltshire, Dan Zhu, and Wolfgang Buermann. 2020. “Climate-Driven Variability and Trends in Plant Productivity over Recent Decades Based on Three Global Products”. Global Bio-geochem. Cycles 34.12, e2020GB006613. doi:10.1029/2020GB006613.

Oksanen, Lauri, Stephen D. Fretwell, Joseph Arruda, and Pekka Niemela. 1981. “Exploitation Ecosystems in Gradients of Primary Productivity”. Am. Nat. 118.2, pp. 240–261. doi:10.1086/283817.

Oksanen, Lauri and Tarja Oksanen. 2000. “The Logic and Realism of the Hypothesis of Exploitation Ecosystems”. Am. Nat. 155, pp. 703–723. doi:10.1086/303354.

Pimm, Stuart L. and John H. Lawton. 1977. “Number of Trophic Levels in Ecological Communities”. Nature 268.5618, pp. 329–331. doi:10.1038/268329a0.

Rip, Jason and Kevin McCann. 2011. “Cross-Ecosystem Differences in Stability and the Principle of Energy Flux”. Ecol. Lett. 14.8, pp. 733–740. doi:10.1111/j.1461-0248.2011.01636.x.

Rosenzweig, Michael L. 1971. “Paradox of Enrichment: Destabilization of Exploitation Ecosystems in Ecological Time”. Science 171.3969, pp. 385–387. doi:10.1126/science.171.3969.385.

Rosenzweig, Michael L. and Robert H. MacArthur. 1963. “Graphical Representation and Stability Conditions of Predator-Prey Interactions”. Am. Nat. 97.895, pp. 209–223. doi:10.1086/282272.

Roy, Shovonlal and J. Chattopadhyay. 2007. “The Stability of Ecosystems: A Brief Overview of the Paradox of Enrichment”. J. Biosci. 32.2, pp. 421–428. doi:10.1007/s12038-007-0040-1.

Schmitz, Oswald J. 2008. “Herbivory from Individuals to Ecosystems”. Annu. Rev. Ecol. Evol. Syst. 39, pp. 133–152. doi:10.1146/annurev.ecolsys.39.110707.173418.

Sharp, Ben R. and Robert J. Whittaker. 2003. “The Irreversible Cattle-Driven Transformation of a Seasonally Flooded Australian Savanna”. J. Biogeogr. 30.5, pp. 783–802. doi:10.1046/j.1365-2699.2003.00840.x.

Shurin, Jonathan B., Elizabeth T. Borer, Eric W. Seabloom, Kurt Anderson, Carol A. Blanchette, Bernardo Broitman, Scott D. Cooper, and Benjamin S. Halpern. 2002. “A Cross-Ecosystem Comparison of the Strength of Trophic Cascades”. Ecol. Lett. 5.6, pp. 785–791. doi:10.1046/j.1461-0248.2002.00381.x.

Su, Haojie, Yuhao Feng, Jianfeng Chen, Jun Chen, Suhui Ma, Jingyun Fang, and Ping Xie. 2021. “Determinants of Trophic Cascade Strength in Freshwater Ecosystems: A Global Analysis”. Ecology 102.7, e03370. doi:10.1002/ecy.3370.

Tan, Zhengxi, Rattan Lal, and Keith D. Wiebe. 2005. “Global Soil Nutrient Depletion and Yield Reduction”. J. Sustain. Agr. 26.1, pp. 123–146. doi:10.1300/J064v26n01_10.

Tilman, David. 1977. “Resource Competition between Plankton Algae: An Experimental and Theoretical Approach”. Ecology 58.2, pp. 338–348. doi:10.2307/1935608.

Vergés, Adriana, Peter D. Steinberg, Mark E. Hay, Alistair G. B. Poore, Alexandra H. Campbell, Enric Ballesteros, Kenneth L. Heck, David J. Booth, Melinda A. Coleman, David A. Feary, Will Figueira, Tim Langlois, Ezequiel M. Marzinelli, Toni Mizerek, Peter J. Mumby, Yohei Nakamura, Moninya Roughan, Erik van Sebille, Alex Sen Gupta, Dan A. Smale, Fiona Tomas, Thomas Wernberg, and Shaun K. Wilson. 2014. “The Tropicalization of Temperate Marine Ecosystems: Climate-Mediated Changes in Herbivory and Community Phase Shifts”. Proc. R. Soc. B: Biol. Sci. 281.1789, p. 20140846. doi:10.1098/rspb.2014.0846.

Volterra, Vito. 1931. “Variations and Fluctuations of the Number of Individuals in Animal Species Living Together.” Anim. ecol., pp. 412–433.

Vuorinen, Katariina E. M., Tarja Oksanen, Lauri Oksanen, Timo Vuorisalo, and James D. M. Speed. 2021. “Why Don’t All Species Overexploit?” Oikos 130.11, pp. 1835–1848. doi:10.1111/oik.08358.

White, Thomas C. R. 2005. Why Does the World Stay Green?: Nutrition and Survival of Plant-eaters. Csiro Publishing.

Yodzis, Peter and Stuart Innes. 1992. “Body Size and Consumer-Resource Dynamics”. Am. Nat. 139.6, pp. 1151–1175. doi:10.1086/285380.

